# 3D printed bioelectronic scaffolds with soft tissue-like stiffness

**DOI:** 10.1101/2024.07.19.604334

**Authors:** Somtochukwu S. Okafor, Jae Park, Tianran Liu, Anna P. Goestenkors, Riley M. Alvarez, Barbara A. Semar, Justin S. Yu, Cayleigh P. O’Hare, Sandra K. Montgomery, Lianna C. Friedman, Alexandra L. Rutz

## Abstract

3D printing is a leading technique for fabricating tissue engineering scaffolds that facilitate native cellular behavior. Engineering scaffolds to possess functional properties like electronic conductivity is the first step towards integrating new technological capabilities like stimulating or monitoring cellular activity beyond the traditionally presented biophysical and biochemical cues. However, these bioelectronic scaffolds have been largely underdeveloped since the majority of electrically conducting materials possess high stiffness values outside the physiological range and that may negatively impact desired cell behavior. Here, we present methods of 3D printing poly(3,4-ethylenedioxythiophene):poly(styrene sulfonate) (PEDOT:PSS) hydrogel scaffolds and provide techniques to achieve stiffness relevant to many soft tissues (<100 kPa). Structures were confirmed as ideal tissue scaffolds by maintaining biostability and promoting high cell viability, appropriate cell morphology, and proliferation. With these findings, we contribute a customizable 3D platform that provides favorable soft cellular microenvironments and envision it to be adaptable to several bioelectronic applications.

## Introduction

In vitro engineering tissues that closely resemble the native physiology are important for developing models for drug discovery and personalized medicine as well as for presenting functional, implantable therapeutics^1,2^. To best support tissue engineering, there has been a shift from 2D to 3D cell culture substrates to best mimic how cells exist in vivo. In fact, many cells require a truly 3D microenvironment in order to assemble into physiologically relevant tissue structures in vitro^2^. One such 3D culture technique includes the use of scaffolds, which are microporous structures that resemble the native extracellular matrix (ECM)^3^. Pores are a key feature of scaffolds and manipulation of their size, shape and distribution has been shown to influence cell behavior including attachment, differentiation, spreading and proliferation^4^. Due to the capability of fine control over pore architecture, 3D printing has become one of the leading strategies for scaffold fabrication^5^. Additionally, this fabrication methodology can generate fully interconnected microporosity that is beneficial for achieving a uniform cell distribution and promoting nutrient and waste transport throughout the scaffold volume^5,6^.

Beyond 3D matrix demands, there have been evolving needs to integrate electronic capabilities to digitally monitor or control cell behavior^7^. Towards this goal, bioelectronic devices such as chopstick electrodes and microelectrode arrays have been employed. Some examples include the use of electrode arrays to record action potentials and field potentials of neurons which are indicative of neuronal function^8,9^, and the use of electrochemical impedance spectroscopy as a direct measurement of cell viability, proliferation, metabolism and death^10–12^. An additional use of these devices, electrical stimulation has been shown to promote cell proliferation and differentiation of several cell types including cardiomyocytes and neurons^13,14^. However, these devices are largely compatible with 2D, monolayer cell culture and have not been adapted for integration with more modern forms of cell culture like 3D scaffolds. Moreover, these devices are often made of metal and silicon which present GPa stiffness. Large deviations by orders of magnitude from the stiffness of the native tissue can lead to non-physiological cell characteristics^15,16^ or even apoptosis^17^. Beyond coarse stiffness adjustments, it is further ideal for many cell types to have substrates that closely match native extracellular matrix (ECM) stiffness to promote appropriate mechano-responses. Differences of as little as a few kPa can have significant impacts on cell behaviors such as changing the metastatic potential of cancer and promoting stem cell differentiation into the desired lineage^18,19^. In order to better benefit from the capabilities bioelectronic devices offer in vitro, there is a need to design electronically conducting materials with soft and 3D characteristics to be integrated with scaffolds or other 3D culture methods.

One approach to integrating technological interfacing into 3D culture is to build scaffolds themselves from functional materials^20^. Hydrogels, a class of materials utilized for their tissue-like properties, are commonly investigated for constructing scaffolds^21^. Of potential electronically conducting materials to fabricate hydrogels, conducting polymers, especially poly(3,4-ethylenedioxythiophene):poly(styrene sulfonate) (PEDOT:PSS), have been widely studied for biointerfacing applications due to its commercial availability, biostability, and biocompatibility^22,23^. Given the aforementioned advantages for control of scaffold properties and engineering cell-material interactions, we sought to fabricate scaffolds by 3D printing PEDOT:PSS hydrogels. In the past few years, several methods have been developed for extrusion-based^24–34^ or light-based^35–37^ 3D printing PEDOT:PSS hydrogels. However, the majority of these studies utilize 3D printing for patterning on substrates resulting in 2D conformable electrodes for implantable or wearable applications rather than for 3D in vitro interfacing. Additionally, it is still challenging to fabricate 3D printed conducting polymer structures which have soft tissue-matching stiffness (1 – 100 kPa) while maintaining good electrical conductivity (>100 S/m). This limitation is likely because high concentrations of PEDOT:PSS often needed for printability or post-processing steps such as annealing for increasing conductivity result in increased modulus.

In this work, we report an approach to engineer conducting, 3D printed PEDOT:PSS hydrogel scaffolds. By utilizing weak gel inks from low concentrations of PEDOT:PSS, and employing embedded 3D printing, millimeter-scale hydrogels were obtained. Additionally post-treatment with solvents or acids influenced hydrogel properties resulting in soft tissue range stiffness (6.20 – 99.8 kPa) with high conductivity up to 1891 S/m. Moreover, we elucidate that the water fraction, conductivity and modulus are strongly correlated with each other and that a balance between properties is required to obtain conditions desirable for biointerfacing. To evaluate biostability, hydrogel conductivity in cell culture conditions was characterized. We identified that hydrogels require an incubation time of a few days before conductivity remains stable over several weeks. Finally, 3D printed PEDOT:PSS hydrogel scaffolds with interconnected porosity were fabricated and supported high cell attachment, viability (>99%), proliferation and spreading with desired morphology. Overall, our results validate that these 3D printed conducting polymer hydrogels can serve as scaffolds by providing a soft, 3D and cytocompatible microenvironment for cells. In the future, these scaffolds show great potential for their integration into bioelectronic devices by electronically connecting these to instrumentation to use as electrodes for monitoring, recording, and control of cell behavior within tissue models.

## Results

### Formulation of PEDOT:PSS weak gel inks for extrusion 3D printing

We and others have previously demonstrated that one approach to achieving printability in hydrogel materials is to utilize a weak gel state as the ink^38–41^. These are mixtures which are after the gel point as determined by the storage modulus (G’ – elastic behavior) greater than the loss modulus (G” – viscous behavior)^38^. Such weak gels (G’∼ 1-100 Pa) can generally be both pushed through fine diameter nozzles yet are shape-maintaining, important for extrusion and achieving defined microarchitectures respectively. We therefore sought to develop similar weak gel inks with the conducting polymer PEDOT:PSS. Hydrogels of PEDOT:PSS have been made from its colloidal dispersions (1.1 – 1.3 wt%) by mixing with ionic liquids (IL) and allowing to gel with time at room or elevated temperatures^42,43^. Ionic liquids induce gelation by performing charge screening of PEDOT and PSS, thereby promoting structural rearrangement of PEDOT and the formation of physical cross-links including through π–π stacking^44^. Our previous work has shown that specific concentrations of ionic liquid (>10 mg/mL) result in mixtures which transition from liquid to gel on a timescale of minutes to hours^42^. We sought to determine if time and IL concentration could be used to obtain a weak gel state as an ink for extrusion 3D printing.

The printability of PEDOT:PSS mixtures containing ionic liquid concentrations of 20 – 100 mg/mL was qualitatively investigated by performing single layer 3D printing tests on glass slides (Figure 1A, Supplementary Figure 1). All mixtures were indeed confirmed to be extrudable. Further classification of printing quality indicated that mixtures of IL concentration ≤ 30 mg/mL did not retain its as-printed shape and spread dramatically on the substrate. Alternatively, mixtures of IL concentrations 40 – 100 mg/mL maintained shape fidelity, although IL concentrations ≥ 80 mg/mL resulted in fragmented printed lines. IL concentrations 40 – 60 mg/mL were therefore selected as inks for ideal printing properties of both shape fidelity and continuous extrusion. Next, the rheological properties of the 40 mg/mL ink were investigated to understand the material behavior under shear stress that enables extrusion. By performing oscillatory time sweeps, the weak gel state was confirmed as the ink transitioned from liquid to gel at ∼20 mins (G’>G”, tan(δ) < 1) (Figure 1B). Within the first hour of gelation, a rapid increase in G’ was observed; however, a more stable and modest growth was obtained after two hours. Printing often occurs over prolonged periods, so it is important to obtain inks that exhibit minimal temporal changes. Thus, inks were used for printing at 2 hours post-preparation for all future experiments (G’_2hr_= 20 - 30 Pa). Furthermore, this PEDOT:PSS ink exhibited shear-thinning properties with viscosity decreasing with increased shear rate (Figure 1C). Additionally, while the ink behaved like a solid at low strains (1%), the material yielded and transitioned to a liquid at higher strains (∼350% for G”>G’ transition) (Figure 1D). We then investigated whether the deformation is permanent or reversible. Reversible deformation after removal of shear, also called shear recovery, is desirable for 3D printing to ensure shape retention after deposition^45^. Upon removal of a high strain (1000%), the ink quickly regained its gel state (G’>G”) (Figure 1E). Approximately 50% of the initial modulus was recovered within one second and full recovery was achieved within 32 mins.

**Figure 1.**
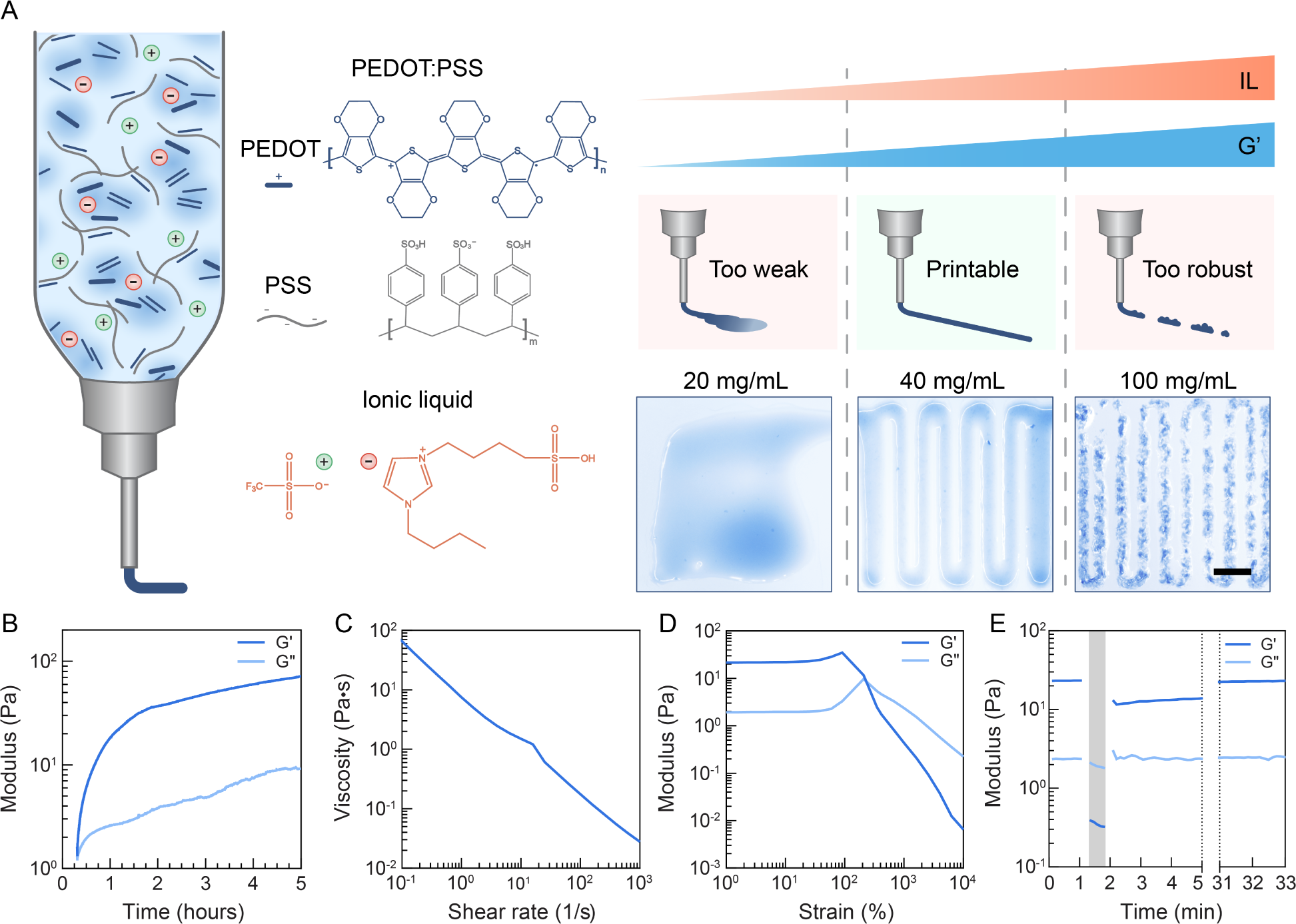
PEDOT:PSS weak gel inks obtained by the manipulation of IL concentration exhibit properties suitable for extrusion 3D printing. A) Schematic of PEDOT:PSS weak gel ink. Ink made by combining PEDOT:PSS and IL gelation agent. By varying IL concentration, inks which are extrudable and produce continuous and defined printed lines are achieved. Scale bar = 2 mm. B-E) Rheological characterization of PEDOT:PSS inks with 40 mg/mL IL showing B) the weak gel state is obtained and that after 2 hours, ink is more stable as assessed by modest changes in G’ (1% strain, 1 rad⋅s^-1^). C) viscosity decreases with increasing shear rate. D) shear yielding from gel to liquid at higher strains (>350%, 1 rad⋅s^-1^). E) material deformation at a high strain (shaded, 1000% strain, 1 rad⋅s^-1^) is recovered when the material is returned to a low strain (unshaded, 1% strain, 1 rad⋅s^-1^).

### 3D printing of multi-layer and free-standing PEDOT:PSS hydrogels

The formulated PEDOT:PSS inks were suitable for fabricating simple structures, such as planar prints. However, the very soft state made printing of more complex and multi-layer objects challenging due to gravity. This capability is crucial for mimicking the native 3D microenvironment for cells through the fabrication of 3D microporous scaffolds, generally ranging of sizes from a few hundred microns to centimeters. Embedded 3D printing can be used to tackle this structural challenge by extruding into a temporary support medium. The support medium possesses rheological properties of shear thinning for ink deposition and shear recovery for physical support until crosslinking is complete^46–48^. Granular hydrogels are particularly suitable as a support medium for hydrogel fabrication due to their gentle removal methods, which do not require high temperatures (>100 °C) or use of harsh solvents that may damage hydrogels^49^.

In this work, agarose and Carbopol® granular hydrogels were investigated since these maintain integrity with the mild temperature processing step (60 °C) that is necessary for PEDOT:PSS gelation (Figure 2A, Supplementary Figure 2). Carbopol® support medium was found to be incompatible with PEDOT:PSS gelation conditions and prohibited its full removal from printed structures (Supplementary Figure 2C). When the PEDOT:PSS ink was extruded into an agarose support medium, the printed structures were observed to diffuse over time (∼1 hour) (Figure 2B, Supplementary Figure 3), which has previously been reported for low viscosity inks^50^. One strategy to eliminate ink diffusion is to quickly crosslink upon entry into the support medium. Our previous work has shown that increasing the concentration of ionic liquid increases the crosslinking rate of PEDOT:PSS hydrogels^42^, and thus the effect of adding IL to the support medium was investigated. Ionic liquid mitigated diffusion and as IL concentration increased, the resolution improved as quantified by a decrease in the strut to nozzle diameter ratio (Figure 2C, Supplementary Figure 3). At an IL concentration in support medium of 70 mg/g, a near perfect ratio of 1.07 was obtained and this was selected for all further studies. To examine potential changes in support medium properties that might have improved printing resolution, rheological characterizations were performed. Shear thinning and shear recovery properties were maintained after addition of IL (Figure 2D, 2E, Supplementary Figure 4). Slight differences observed in the support medium when containing ionic liquid were an increase in viscosity and G’ as well as a faster recovery. Storage modulus recovered ∼90% immediately after removal of a high strain and full recovery within 2 minutes, as compared to the support medium without IL which displayed ∼65% immediate recovery and full recovery in greater than 5 minutes. This fast shear recovery of the support medium is important to trap the ink in place during printing^49^. In all, the increased ink crosslinking rate^42^ as well as the increase in viscosity^49^, and shear recovery speed of the support medium are likely contributing to the improved resolution.

**Figure 2.**
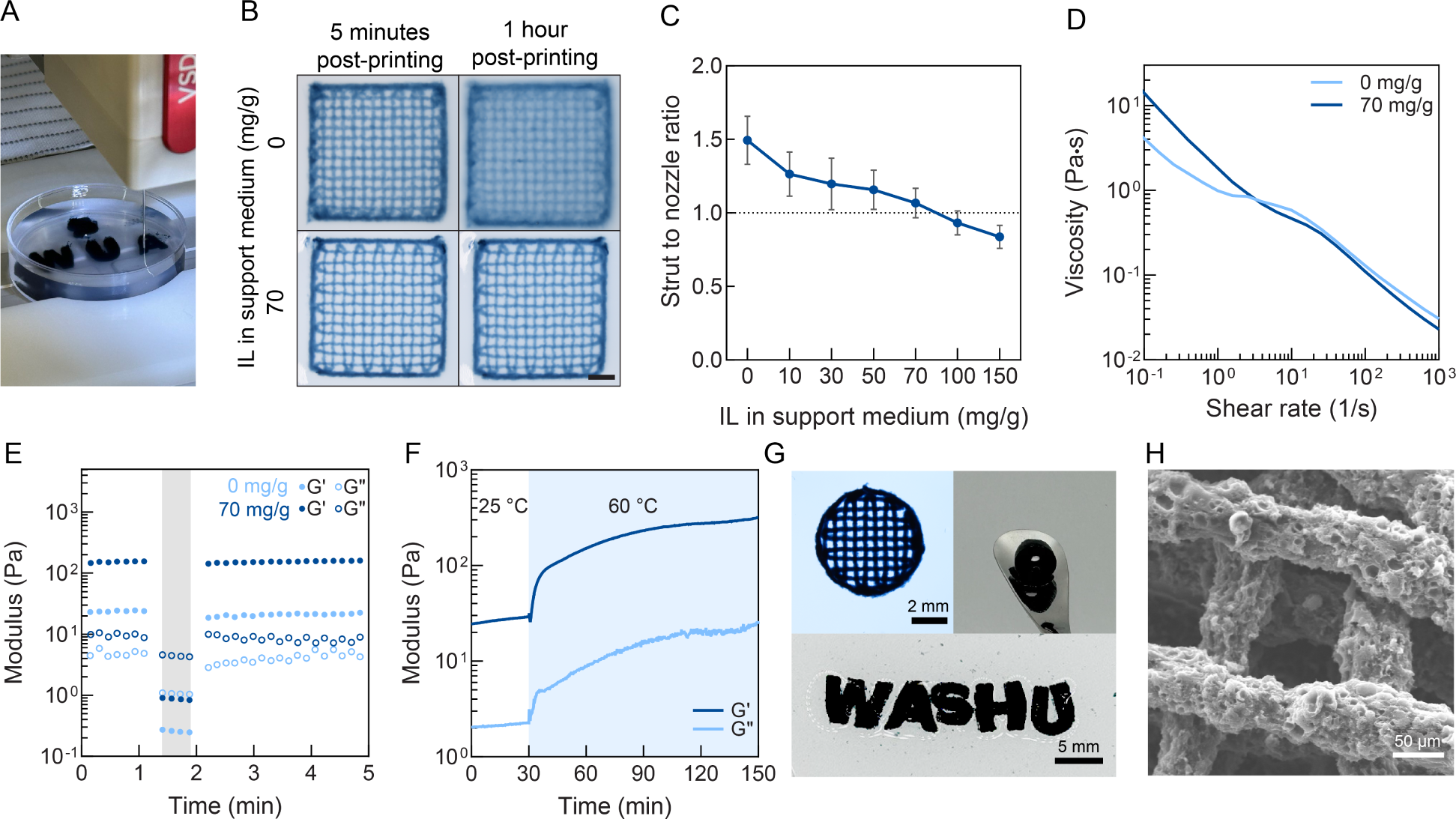
Embedded 3D printing for the fabrication of free-standing, multi-layer PEDOT:PSS hydrogels. A) Representative image of embedded 3D printing of PEDOT:PSS into agarose granular gel support medium. B) Images of ink diffusion into support medium with time. C) Resolution of printed structure versus ionic liquid concentration in support medium measured at 1 hour after printing. The addition of IL improves the printing resolution, dotted line indicates a perfect strut size to nozzle ratio of 1. N≥10. Mean and standard deviation presented. D-E) Rheological characterization of agarose support medium with and without IL addition showing that D) viscosity decreases with increasing shear rate for both conditions and E) material deformation from a high strain (shaded, 100% strain, 1 rad⋅s^-1^) recovers at a low strain (unshaded, 0.1% strain, 1 rad⋅s^-1^). F) Rheological characterization of PEDOT:PSS inks with 40 mg/mL IL at room temperature (unshaded, 25°C) and elevated temperatures (shaded, 60°C) indicating the faster gelation kinetics at higher temperatures (1% strain, 1 rad⋅s^-1^). G) Free-standing, millimeter-scale PEDOT:PSS hydrogels. Cylinder on spatula is ∼8 mm outer diameter and ∼3 mm inner diameter. H) Representative SEM image of printed hydrogels showing clear pored indicating successful support medium removal.

After printing was completed, prints were heated overnight at 60 °C to expedite further crosslinking from the weak gel state to hydrogels expected to be aqueous stable and robust for handling. The effect of raising the temperature is evident from rheology; gelation rate rapidly increased as inks transition from 25 °C to 60 °C (Figure 2F). After the heating period, the support medium was removed to obtain free-standing PEDOT:PSS hydrogels by washing the printed structures in deionized water at 90 °C to melt the agarose. Through this method, free-standing PEDOT:PSS hydrogels of various sizes and shapes were created, including scaffolds for cell culture to be used in later experiments (Figure 2G). The successful removal of the support medium was confirmed by visualizing cleared scaffold pores using scanning electron microscopy (SEM) (Figure 2H). Washed hydrogels were then immersed in de-ionized water to allow for equilibrium swelling. The time to reach equilibrium swelling is important to determine in order to perform characterization of hydrogels in the fully swollen state. Swelling was characterized by measuring dimensional changes in the *xy* and *z* directions. Equilibrium swelling was reached within 24 hours with no significant changes for the rest of period (Supplementary Figure 5). These measurements also indicated an aqueous stable PEDOT:PSS hydrogel network was formed. Swelling was nearly isotropic (*xy*/*z* =1.05 ± 0.08) and this is desirable since it helps maintain the relative dimensions of the print design after fabrication.

### Post-fabrication treatment with acids or organic solvents modulates properties of PEDOT:PSS hydrogels

To expand utility of bioelectronic scaffolds across cell types, it is important to manipulate properties to best match characteristics of the ECM of the target tissue (i.e. physiomimetic models). Of the spectrum of tissues, many will possess stiffness in the softest regime of ∼1-100 kPa, including nervous, muscle, and adipose tissues as well as most internal organs^15^. In typical hydrogels used for tissue engineering, stiffness is often altered by changing the polymer concentration or the degree of crosslinking. Crosslinking in PEDOT:PSS hydrogels is achieved through enrichment of PEDOT, resulting in increased π-π bonds, and thereby stiffness. Particularly, in PEDOT:PSS thin films (1 nm – 1 μm thickness), methods to elicit PEDOT enrichment have been significantly investigated for improving electronic conductivity. One method is to treat films post-fabrication with acids or organic solvents^51–54^. These treatments act by inducing charge screening of PEDOT and PSS and lead to loss of PSS from the material to the surrounding solvent. While established in thin films, these have been limitedly studied for hydrogels specifically^26,27^. Furthermore, how these treatments can be used to modulate the stiffness of PEDOT:PSS has not yet been reported. Given that manipulation of cross-linking in turn impacts several other properties of hydrogels, these treatments were anticipated to additionally change swelling and conductivity. In this work, we selected sulfuric acid (18.0 M), acetic acid (10.4 M), ethanol and dimethyl sulfoxide (DMSO) as example treatments and investigated their effect on these properties of the 3D printed PEDOT:PSS hydrogels.

The majority of previously reported methods for treating PEDOT:PSS utilize high temperatures (>100 °C) that may completely dehydrate or damage hydrogels. Thus, we adapted the methods to be compatible by performing at room temperature. All post-treatments were carried out by immersing the hydrogels in the solvents or acids overnight followed by rigorous washing with water (Figure 3A). To characterize the effect of treatment on stiffness, the storage modulus (G’) was measured. The G’ of the hydrogels after full fabrication (complete gelation and support medium removal) were 2.90 ± 0.914 kPa (Supplementary Figure 6A-B, Supplementary Table 1). Significant changes in G’ were observed only for acid-treated hydrogels (P<0.0001; Supplementary Figure 6A-B, Supplementary Table 1). Elastic modulus was calculated assuming a Poisson’s ratio of 0.5. Collectively, these various treatments covered the range of elastic modulus relevant for soft tissues, from 6.20 – 99.8 kPa (Figure 3B). Notably, all samples had shrunk after post-treatment, with the most substantial shrinkage relative to as-fabricated observed in the acid-treated hydrogels (Figure 3B, Supplementary Figure 6C). Shrinking was nearly isotropic across all samples (*xy/z* = 0.94 – 1.16) which benefits the maintenance of the printed design relative to itself.

**Figure 3.**
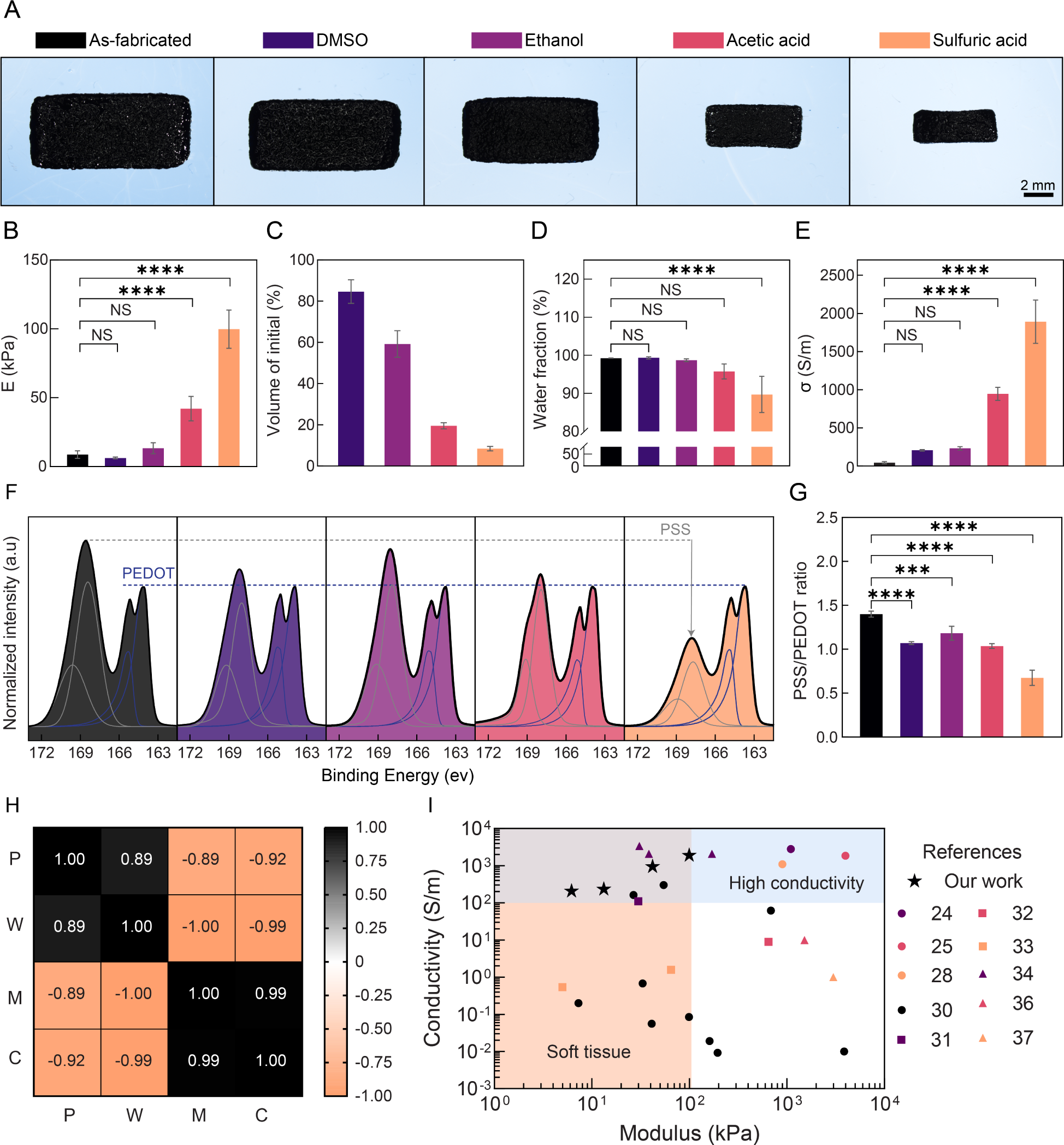
Post-treatment of PEDOT:PSS hydrogels results in tunable material properties. A) Representative images of samples after post-treatment. B-E) Post-treatment with solvents or acids influences the B) elastic modulus, C) volumetric swelling (percentage relative to as-fabricated hydrogels, without treatment), D) water fraction and E) electronic conductivity of hydrogels. F-G) X-ray S 2p photoelectron spectra of treated and as-fabricated hydrogels indicates a decrease in PSS after all treatments. The de-convoluted profiles were fitted with asymmetric functions. Mean and standard deviation presented. Quantification of PSS/PEDOT ratio calculated from a comparison of the area under the curves of the XPS fits. N≥4. One-way analysis of variance (ANOVA) and Dunnett’s multiple comparison test performed. ****P≤0.0001, ***P≤0.001, **P≤0.01, *P≤0.05, Non-Significant (NS) P>0.05. H) Pearson’s correlation heatmap showing the correlation between PSS/PEDOT ratio (P), water fraction (W), elastic modulus (M) and conductivity (C). I) Survey of recent literature that report modulus and conductivity of 3D printed PEDOT:PSS hydrogels. High conductivity often leads to stiffness values outside the physiological range. The present study joins very few others able to meet satisfactory conductivity with elastic moduli representing soft tissues.

Following these trends in swelling differences, slight changes in the water fraction of hydrogels were also measured. After treatment, a small decrease in mean water fraction was observed for acetic and sulfuric acid treatments, with a significant change for sulfuric acid treated gels after correcting for multiple comparisons (P<0.0001, Figure 3D, Supplementary Table 1). Despite these changes, the water fraction of the gels after treatment remained high (89.7-99.4%), indicative of maintenance of the hydrogel state (typically ≥90%). Such high water content is beneficial as the polymer network can absorb media components including nutrients which is important for maintaining cell viability and proliferation throughout the scaffold volume. Finally, the conductivity of the printed gels increased after all treatments, with significant increases observed for acid-treated hydrogels (as high as a 40-fold increase to 1891 S/m; Figure 3E). Increasing conductivity may be advantageous for scaffolds as cell attachment and spreading of certain cell types on substrates have been shown to increase with conductivity^55^. Additionally, higher conductivity would be expected to yield better performance for bioelectronic devices based on these scaffolds. Here, we explored four common post-treatments, but we envision others can also be used (e.g. isopropanol, methanol, ethylene glycol). Beyond solvent type, additional variables such as concentration and time of exposure may lead to even further tailoring of properties. To support this idea, we found that when the molarity of the acids was decreased, conductivity and modulus decreased while water fraction increased (Supplementary Figure 7).

Altogether, these treatments generally increased modulus, increased conductivity, and decreased swelling, with the acids being most effective. To confirm the cause of these trends, we performed X-ray photoelectron spectroscopy (XPS). In the sulfur 2p orbital spectra (S 2p), the binding energy peaks within 161–165 eV represent the thiophenes in PEDOT, while peaks within 166–171 eV correspond to sulfonate groups in PSS^56^ (Figure 3F). A significant decrease in the calculated PSS/PEDOT ratio was observed after all treatments when compared to the untreated hydrogel (P≤ 0.0001, Figure 3G, Supplementary Table 1). A decrease in the PSS/PEDOT ratio confirmed PSS removal and is likely indicative of larger PEDOT-rich regions and increased π-π bonding among PEDOT chains^56^. This more tightly crosslinked network is anticipated to contribute to the increased modulus and higher conductivity generally observed after treatment. Beyond increased crosslinking, there could be additional factors responsible. For example, the loss of hydrophilic network components (PSS) relative to hydrophobic components (PEDOT) likely contributes to the decrease in water fraction, and water content itself influences matrix viscoelasticity.

It is important to note that the conductivity, modulus, water fraction and PSS/PEDOT ratio were strongly correlated with each other (Pearson’s correlation coefficient ≥ ±0.89, Figure 3H). This means that as one seeks to optimize one property for the intended tissue model, these other factors are anticipated to change as a result. For example, one may desire higher conductivities and thus perform additional processing, but this target property may come at a risk of changing stiffness and water fraction beyond the desired range for in vitro biointerfacing. Our work joins very few others able to achieve a satisfactory balance of modulus and conductivity, and further, is able to do so over a range of moduli (Figure 3I).

### 3D printed PEDOT:PSS hydrogel scaffolds for in vitro biointerfacing

Majority of in vitro biointerfacing applications will require studies to be carried out over periods ranging from days to months. Thus, it is important to characterize the stability of material over time to determine suitability for use. A few works have shown PEDOT:PSS hydrogels to be stable (mass, conductivity, swelling) in deionized water and phosphate buffered saline (PBS) over extended periods at room temperature^42,57^. Such methods may not be sufficient for evaluation since there are substantial differences in incubation media composition (salts, amino acids, and vitamins) as well as temperature (37 °C) which may potentially induce changes in the material.

In this work, the stability of PEDOT:PSS hydrogels incubated in Dulbecco’s Modified Eagle Medium (DMEM) was investigated for a duration of 28 days (37 °C, 5% CO_2_). Throughout the duration of the study, the hydrogels showed no visible fragmentation and no differences in ease of handling compared to those prior (Figure 4A, Supplementary Figure 8A). The mass of the hydrogels was also measured over time to examine for any potential polymer network loss. No significant changes were observed for both conditions throughout the duration of the study (Figure 4B). Given that bioelectronic scaffolds will likely be utilized for their functional property of electronic conductivity as either a matrix cue or as part of a device, it is important to characterize this property over a cell culture period^58^. After initial incubation in DMEM, conductivity decreased ∼30-40% with these maximum decreases occurring at 3 and 7 days for acetic acid and DMSO treated samples, respectively (Figure 4C). Stable conductivity values were observed for the remaining period of three weeks for both conditions (>200 S/m). Our results show similar trends with Guex et al.^59^ and Marzocchi et al.^60^, which, to the best of our knowledge, are the only other works that have reported the conductivity of PEDOT:PSS over time in cell culture conditions.

**Figure 4.**
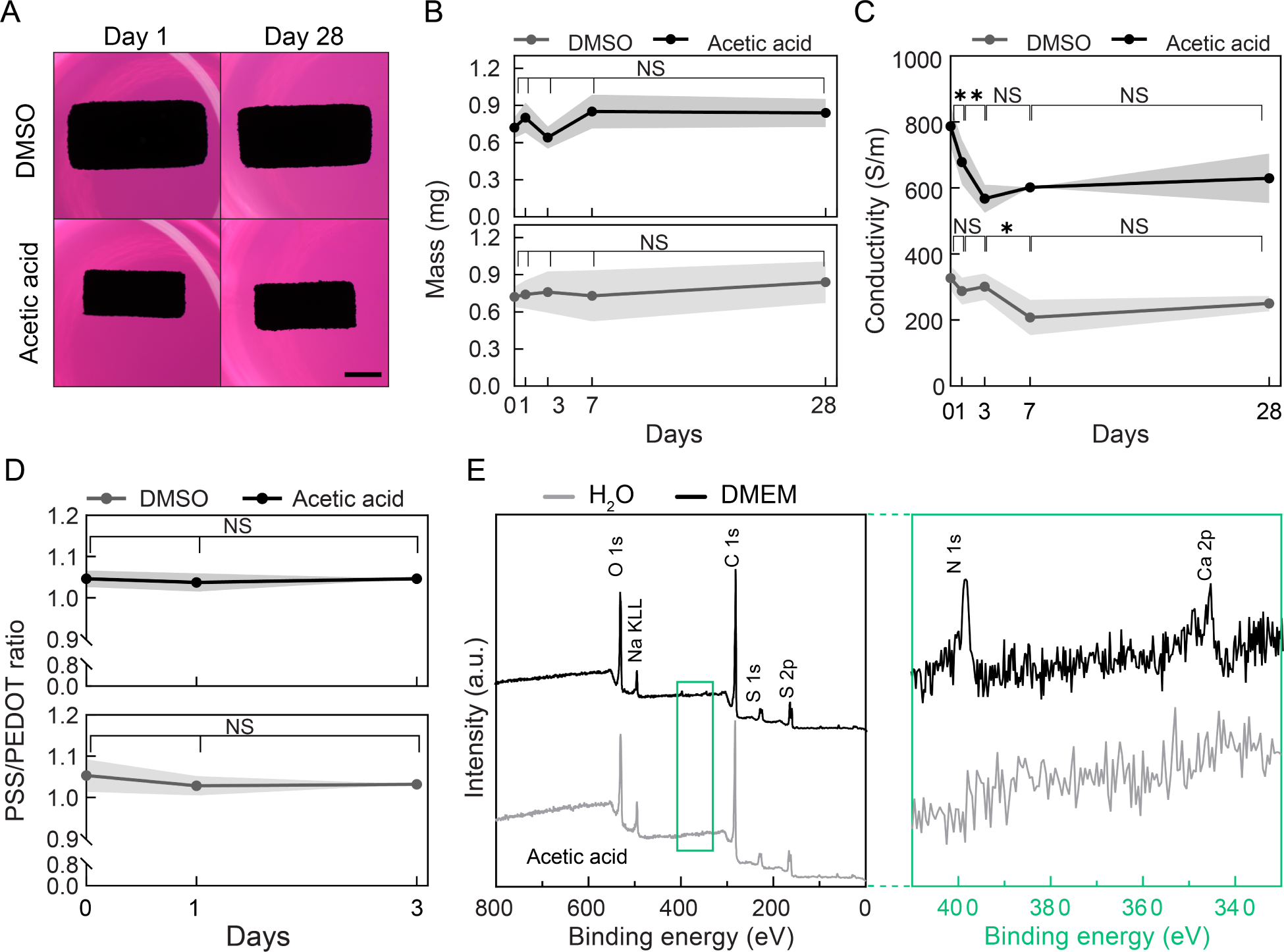
PEDOT:PSS hydrogels reach stable conductivity within days during incubation in cell culture conditions. A) Representative images of hydrogels post-treated with DMSO and acetic acid, scale bar = 2 mm. Hydrogels show no visible fragmentation or breakdown over 28 days for both conditions. B) Dry mass of hydrogels over time. No significant changes in mass for both conditions. C) Conductivity of PEDOT:PSS hydrogels decreased during initial days of incubation but stabilized for both conditions after seven days. D) No significant differences in the PSS/PEDOT ratio for first three days in DMEM. One-way analysis of variance (ANOVA) and Šidák multiple comparison test performed. Non-Significant (NS) P>0.05. E) XPS of acetic acid treated samples before (in deionized water) and after incubating in DMEM showing presence of additional peaks. Green boxes highlight the binding energy range covering nitrogen and calcium. Mean and standard deviations presented. N≥3.

To probe why conductivity decreased in the initial days after DMEM incubation, XPS was performed to determine if there were changes in chemical composition. Hydrogels incubated in DMEM were first compared to those incubated in PBS. No significant differences in the PSS/PEDOT ratio were observed during the incubation among days 0, 1, and 3 (Figure 4D). This signifies that the cell culture conditions were not affecting the content of PEDOT and PSS, which aligned with the result of no significant mass change. When compared to post-treated PEDOT:PSS hydrogels swollen in water, media exposed samples displayed new peaks indicative of presence of nitrogen and calcium (Figure 8E, Supplementary Figure 8B). Prior to analysis, these hydrogels had been thoroughly washed in deionized water to remove absorbed media. Our results suggest that PEDOT:PSS is adsorbing calcium ions and nitrogen-containing compounds (e.g. amino acids), and such adsorption may result in structural changes of the polymer network thereby influencing conductivity. These characterizations provide insights that are beneficial for future works integrating PEDOT:PSS based films or hydrogels into in vitro systems as these results especially highlight the likely need to equilibrate materials prior to use as part of devices. Beyond cell culture, similar changes may occur when implanted PEDOT:PSS is exposed to interstitial fluid, blood, or other biological fluids.

3D printed PEDOT:PSS hydrogel scaffolds were next investigated for their potential to support populations of human primary cells in vitro. Samples were disinfected for cell culture and pre-conditioned in fetal bovine serum which has been shown to improve cell attachment for PEDOT:PSS hydrogels^42^. After seeding normal human dermal fibroblasts, both conditions supported high cell attachment (>74% of seeded cells, Supplementary Figure 9A). Cells displayed a spread morphology characteristic of fibroblasts (sphericity ∼ 0.39, Figure 5A, Supplementary Figure 9B). Additionally, cells maintained high viabilities (>99%) for the seven days studied (Figure 5A-B, Supplementary Figure 10). This result gave us confidence that the washing steps employed are effective at removing cytotoxic ionic liquid and treatment solutions. Cells wrapped around scaffold struts and proliferated over the culture period (Figure 5A, C).

**Figure 5:**
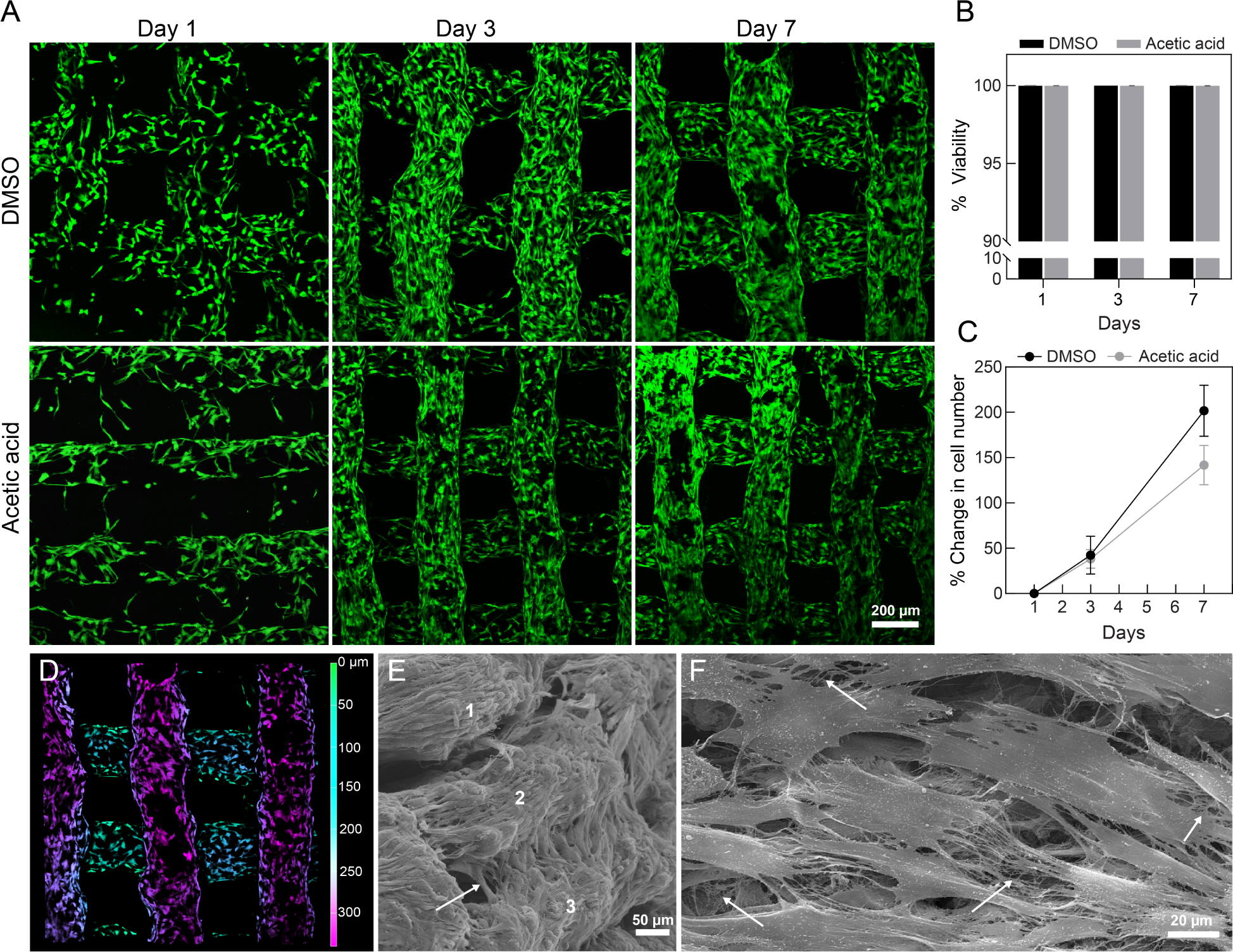
3D printed PEDOT:PSS hydrogel scaffolds support cell attachment and proliferation of human dermal fibroblasts for at least 7 days. A) Representative fluorescence microscopy maximum intensity projections of Z stacks of human dermal fibroblasts on 3D printed PEDOT:PSS hydrogel scaffolds after Live/Dead™ staining (Calcein AM, green; ethidium homodimer-1, magenta). B) Quantification of cell viability from Live/Dead™ staining. C) Quantification of cell number over time. D) 3D depth coded images of maximum intensity projections of Z stacks for cells after one day on DMSO treated scaffolds (>300 μm thickness). E-F) Scanning electron microscopy images of seeded cells on DMSO treated scaffolds after seven days showing E) cells spreading across multiple layers (arrows) with layers identified as numbers 1-3 and F) extracellular matrix deposition by cells on scaffolds. Mean and standard deviation presented. N=4.

Moreover, cells penetrated through multiple layers of the scaffold (Figure 5D-E, Supplementary Figure 9C), highlighting the benefit of the interconnected microporosity. Additionally, fibrous and mesh-like structures resembling deposited extracellular matrix (ECM) were observed at seven days (Figure 5E, Supplementary Figure 9D). This is consistent with other works as fibroblasts particularly in 3D cultures have been shown to deposit ECM as early as one week post seeding^61^. ECM deposition is a key function of dermal fibroblasts to maintain structural integrity of skin and contribute to wound healing. These findings validate that these 3D printed PEDOT:PSS hydrogels can serve as tissue engineering scaffolds and would likely serve as suitable cell culture substrates in the context of 3D bioelectronic devices.

## Discussion

In this work, we report methods to present the conducting polymer PEDOT:PSS as a 3D printed biomaterial scaffold. First, we developed weak gel inks of the appropriate rheological properties (G’∼20 Pa) for extrusion 3D printing. By using these weak gel inks, and by optimizing embedded 3D printing methods, soft, PEDOT:PSS hydrogels (∼8 kPa, >99% water content) were fabricated. Additionally, through post-treatment with acids and organic solvents, hydrogels with high water content (>89%), high conductivities (207 – 1897 S/m) and tissue-range stiffness (∼6-100 kPa) were obtained.

Our results indicate that the amount of PSS removal from these treatments contributed to these changes in material properties. Beyond the effect of the treatments on the individual properties, we also show that the water content, conductivity, modulus and PSS/PEDOT ratio are strongly correlated, indicating that a balance in properties must be considered for improved biointerfacing and device integration. By characterizing the conductivity of hydrogels in cell culture conditions, we determined that a short initial incubation period (3 – 7 days) was required for equilibration of electronic properties and after which, stability and high conductivity were maintained (≥3 weeks, >200 S/m). Further characterization on the chemical composition of the hydrogels after media incubation suggested that media species adsorption rather than changes in the PEDOT and PSS content is likely behind the initial changes in electrical properties. These findings highlight the need for characterization of functional properties of materials in in vitro conditions beyond the more traditional physical characterizations (e.g. mass, dimensions) performed.

By taking advantage of additive manufacturing, PEDOT:PSS hydrogel scaffolds with interconnected porosity were fabricated. The scaffolds supported high viability (>99%) of human primary cells, while providing desired 3D microenvironment for cells to proliferate across the scaffold volume (>500 µm) and produce extracellular matrix. Due to the capabilities 3D printing offers, the dimensions, structure, and porosity of PEDOT:PSS scaffolds can all be customized. For example, multi-scale and controlled gradient porosities^62^ are being actively explored as a technique for improving cell infiltration and better mimicking the native extracellular matrix. This technology can also be developed to be compatible with multi-material fabrication for printing electronic materials alongside biomaterials and cell-laden hydrogels for biointegrated devices.

In the future, these bioelectronic scaffolds can be further developed as devices capable of simultaneously providing an ideal cellular microenvironment as well as having electronic capabilities of recording, stimulation or impedance monitoring. While human tissue models represent most potential uses, other species such as plant cells may also benefit^63^. Beyond these functional cell culture substrates for engineering tissue in vitro, these scaffolds could also serve as host vehicles for implantable cell therapeutics^64^. Electronic scaffolds can provide the stimulus for electrically stimulated drug release^65^ or as niches for cells that serve as biological bridges between technology and host tissue^66^. These applications are part of a boarder class of “biohybrid devices” fusing cells and technology to achieve new properties and function in living technology.

## Methods

### Preparation of PEDOT:PSS 3D printable ink

Colloidal aqueous dispersions of PEDOT:PSS (Clevios PH 1000, 1.0 – 1.3 wt%) were acquired from Heraeus. Ionic liquid (IL), 4-(3-butyl-1-imidazolio)-1-butanesulfonic acid triflate (Santa Cruz Biotechnology) was used as a gelation agent at concentrations 20 – 100 mg/mL. PEDOT:PSS was filtered with 0.45 μm polytetrafluoroethylene (PTFE) syringe filters to remove large aggregates. Inks were prepared by vortex mixing the PEDOT:PSS aqueous dispersion with IL for 30 seconds and then stirring for 15 minutes at 700 RPM. Ink age is determined by the time from which PEDOT:PSS was added to IL.

### 3D printing of PEDOT:PSS hydrogels

All hydrogels were fabricated using a R-GEN 100 3D Bioprinter (RegenHu, Switzerland) outfitted with a volumetric printhead (mechanically driven piston extrusion). Inks were loaded into a Hamilton syringe customized for the R-GEN 100 printer, with a 26-gauge needle with an inner diameter 260 μm, length 25.4 mm (SAI infusion technologies). All print designs were created using Shaper (RegenHu, Switzerland). Printability assessment of inks with varying IL concentrations was done by extruding PEDOT:PSS inks (1 layer, 10 x 10 mm, 500 μm spacing, 10 mm/s, 25 °C) on clean glass slides. Brightfield images were taken using a stereoscope (Nikon SMZ1270).

### Support medium preparation

Agarose granular hydrogels were prepared by autoclaving agarose (0.5% w/v, Sigma-Aldrich) dispersed in deionized water and then cooling autoclaved agarose to room temperature under constant shear of 700 rpm^50^. Note for optimal results, we found that the agarose volume should be less than ∼40% of the media jar volume used for autoclaving (e.g. 200 mL in 500 mL media jar). To prevent ink diffusion in support medium, ionic liquid (IL), 4-(3-butyl-1-imidazolio)-1-butanesulfonic acid triflate (AmBeed) was mixed with the agarose granular hydrogel at varying concentrations (0 -150 mg IL per g support medium) by vortex mixing for 15 minutes and then allowed to sit overnight. Prior to printing, the support medium was centrifuged to remove air followed by a brief and low force vortex (∼10 seconds) to resuspend support medium without introducing air bubbles. Agarose support medium (granular gel + ionic liquid) was stored in centrifuge tubes at room temperature until further use.

### Embedded 3D printing of hydrogels

For the fabrication of 3D structures, PEDOT:PSS inks (40 mg/mL IL, ink age = 2 hours) were loaded into the Hamilton syringe and desired shapes were extruded into the support medium loaded into a 35 mm petri dish (26-gauge nozzle, 3 mm/s, 25 °C). Non-porous shapes were fabricated to be dense (100% infill) by having a smaller slicing than the nozzle size (e.g. a 30-gauge slicing was done for a 26-gauge nozzle). Non-porous structures were generally used for hydrogel material property characterization, while porous (described later) were used for cell-material interfacing experiments. After printing, hydrogels were transferred to a container (polypropylene jar, Nalgene®, 60 mL). Deionized water was added to the bottom of the container to maintain local humidity and prevent significant water evaporation of the support medium. The containers were then incubated at 60 °C for 16 hours for further crosslinking. After oven incubation, hydrogels were washed in deionized water at 90 °C for 1 hour to melt the agarose.

### Hydrogel washing and equilibrium swelling

Once support medium was removed, washed hydrogels were incubated in 10 mL of deionized water to allow for equilibrium swelling. For swelling experiments, non-porous PEDOT:PSS hydrogels (6 mm length, 6 mm width, 2 mm thickness) were printed and measured over time in water. To measure hydrogel dimensions, images of the top (length, width, *XY*) and side (thickness, *Z*) were taken using a stereoscope (Nikon SMZ1270) and measurements were performed with NIS-Elements (Nikon). To determine water fraction of hydrogels, samples were weighed in tared microtubes to obtain wet mass. Then, hydrogels were frozen at -20 °C, lyophilized, and finally weighed to obtain dry mass. The water fraction was calculated using the following equation:

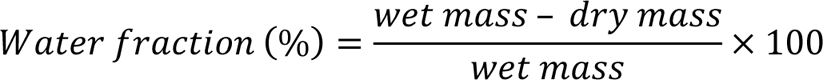

### 3D printed hydrogel post-treatment

3D printed PEDOT:PSS hydrogels were immersed in 4-5 mL of dimethyl sulfoxide (DMSO), ethanol, 10.4 M acetic acid solution and 18.0 M sulfuric acid (18.0 M) overnight. After incubation, hydrogels were washed using exchanges of 10 mL of deionized water each as follows: at least four exchanges every 15 minutes for the first one hour followed by one overnight water exchange. Samples incubated in acidic solutions underwent more rigorous washing with 5 minute water exchanges to bring pH to ∼5 before the above washes were performed.

### Rheological characterization

All rheological characterizations were performed using a TA Instruments HR-20 rheometer. All ink measurements were taken with a cone-plate fixture (stainless steel plate of 40 mm diameter and 1.98889° cone geometry). A solvent trap was used to prevent water evaporation. Viscosity of PEDOT:PSS-IL mixtures was investigated by performing rotational flow sweeps at shear rates of 10^-1^ 1/s – 10^3^ 1/s at 25 °C. Oscillatory sweeps were performed for various evaluations including gelation kinetics of inks as well as shear thinning and shear recovery of both inks and support medium. Unless otherwise noted, oscillatory sweeps for inks were conducted at 25 °C, 1% strain, and 1 rad⋅s^−1^ angular frequency while oscillatory sweeps for support medium were conducted at 25 °C, 0.1% strain, and 1 rad s^−1^ angular frequency.

All printed hydrogels were biopsy punched and characterized using an 8 mm parallel-plate fixture. Stiffness of as-fabricated and post-treated gels was determined using oscillatory stain sweeps at 25 °C and 10 rad⋅s^−1^ angular frequency. A 0.05 N pre-load was applied to determine sufficient contact before measurements on all samples. The region from 0.01 to 0.1 % strain was determined as the linear region. The storage modulus (G) was determined by averaging the values over this range. Elastic modulus (E) was determined using the assumed Poisson’s ratio (*v* =0.5) by the following equation:

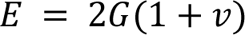

### Electrical characterization

The conductivity of PEDOT:PSS hydrogels was measured at room temperature using a four-point probe (Ossila, UK) with rounded tips for preventing damage to hydrogels. Excess water from hydrogels was removed by dabbing with Kimwipes™. Probes were positioned to be centered on the gel and the stage supporting the hydrogel was slowly raised until all four probes had contacted the surface of the hydrogel. Following manufacturer recommendations, the stage was further raised to confirm sufficient contact between probes and hydrogel. Conductivity was obtained from the Ossila Sheet Resistance Software. Conductivity values presented were taken from a mean of 50 recordings per measurement per sample.

### Scanning electron microscopy (SEM)

Scaffolds were imaged using a Zeiss EVO 10 scanning electron microscope to visualize their morphologies. All samples were crosslinked with 2.5 v/v% glutaraldehyde and 3 wt% sucrose in PBS for 1-2 hours at room temperature. Samples without cells were exchanged in a graded series of ethanol, then completely dehydrated using a critical point drier (K850, Electron Microscopy Sciences). To perform SEM imaging, the dried samples were mounted to stubs using carbon tape. Samples seeded with cells were returned to the physiological osmolarity after initial fixation, and further crosslinked overnight using 5 wt% ethylene glycol diglycidyl ether in pH 9 PBS. Post fixation was performed by treating the samples with 1% osmium tetroxide in 0.1 M sodium cacodylate buffer for 1 hour. Following serial exchange to ethanol, the samples were critical point dried using a Leica EM CPD300 Critical Point Drier. Carbon tape and silver paste were used to mount the samples, which were then sputtered with 10 nm carbon and 12 nm iridium. All SEM images were captured at 20 kV with a working distance ranging from 9 to 20 mm.

### X-ray photoelectron spectroscopy

XPS spectra were obtained using an X-ray photoelectron spectrometer (PHI 5000 VersaProbe II, Physical Electronics), and the hydrogel samples were lyophilized before characterization. The survey spectra for identifying the chemical elements in the samples were recorded with a binding energy step resolution of 0.5 eV. To investigate the chemical properties of PEDOT, the S 2p spectra were recorded with a binding energy step of 0.05 eV. Spectra peaks were calibrated by referencing the binding energy of C 1s (284.8 eV). The XPS spectra of S 2p were fitted using an asymmetric line shape. The areal ratio of the two doublet peaks of S 2p_1/2_ and S 2p_3/2_ was fixed at 1:2, with the binding energy splitting of each doublet peak being 1.2 eV. The PSS-to-PEDOT ratio was calculated using the S 2p doublet peak area corresponding to PEDOT and PSS.

*Assessment of 3D printed PEDOT:PSS hydrogel stability in cell culture conditions* Printed PEDOT:PSS hydrogels were post-treated with DMSO and 10.4 M acetic acid, respectively. Following post-treatment, samples were transferred to 12 well-plates and disinfected by incubating in 70% ethanol (5 mL) overnight at room temperature in a biosafety cabinet. Hydrogels were then washed in sterile 1X phosphate-buffered saline (PBS) with 1% penicillin–streptomycin (Pen/Strep; 10,000 U mL^-1^ penicillin and 10,000 μg/mL streptomycin). Samples were stored in sterile PBS+Pen/Strep (5 mL) at 4°C until study (at least overnight). All samples were handled aseptically in a biosafety cabinet during preparation and for the duration of the study.

To begin assessment, hydrogels were each incubated in 4 mL of Dulbecco’s Modified Eagle Medium (DMEM) (Corning) containing 1% Pen/Strep and then transferred to a cell culture incubator at 37 °C, 95% relatively humidity, and 5% CO2. Hydrogel mass, conductivity, and dimensions were measured as previously described.

### In vitro cell studies

#### Scaffold preparation

For in vitro assessment, cylindrical scaffolds of 6 mm diameter, ∼2 mm thickness (8 layers) and 150 - 300 μm pores with 90° advancing angles were fabricated. For better structural robustness, scaffolds were printed with a bounding perimeter. As described above, scaffolds were post-treated, disinfected in 70% ethanol and washed in PBS+Pen/Strep. Afterwards, samples were transferred to sterile 12-well plates coated with 1 wt% agarose which provides a non-cell-adhesive surface under the scaffold, and then stored in PBS+Pen/Strep overnight at 4 °C. Scaffolds were preconditioned for cell culture by incubating in 1 mL of fetal bovine serum (FBS) at 37 °C overnight and then washed in PBS+Pen/Strep for 15 minutes to rinse the FBS.

#### Cell seeding

Normal adult human dermal fibroblasts (NHDF; Lonza CC2511) were cultured according to supplier instructions and used at passage 5. Immediately prior to cell seeding, scaffolds were dried for ∼45 minutes in a biosafety cabinet in order to remove water held within the micropores and allow the scaffolds to be able wick up the applied cell suspension. The cell suspension (10-15 μL) was deposited drop-wise on dried scaffolds to target a cell density of 45,000 cells/cm^2^ and then transferred to the incubator for 30 minutes to allow for cell attachment. Four mL of cell culture media was then added to each well containing scaffolds and cell culture media was changed every other day.

#### Cell viability and sphericity analysis

Cell-seeded scaffolds were evaluated for cell viability and attachment through fluorescence staining. The staining solution contained 2 µM Calcein AM, 8 µM of ethidium homodimer-1, and 10 µg/mL Hoechst 33342 (BD) in PBS. Each scaffold was transferred to a confocal glass bottom dish and stained with the staining solution for at least 30 minutes. Imaging was performed with a Leica SP8 Lightning confocal microscope. Four regions of interest were imaged per scaffold.

The images were processed using the Leica LAS X 3D Analysis suite and cell viability was obtained by determining total number of all cells and of dead cells and subtracting the two to find the total number of live cells.

Total cells were determined by the total number of nuclei (Hoechst) and dead cells by the number of red stained nuclei (ethidium homodimer-1).

Cell sphericity on one day after seeding was quantified using the in-built sphericity feature in the LAS X 3D Analysis suite on the Calcein AM channel (cell body) after image processing.

#### Cell proliferation assay

Cell-seeded scaffolds were collected in tared microtubes and frozen at -20°C at select time points until time of the assay. Aliquots of the cell seeding suspension were also collected. All samples were then lyophilized and digested in Proteinase K solution overnight at 60 °C. The Picogreen assay was then run to quantify DNA content on scaffolds using the manufacturer’s instructions. Cell attachment efficiency (%) was determined by dividing the live cells present after one day by the cell number from the cell seeding suspension and multiplying by 100.

### Literature search for comparison literature

Literature search on 3D printed PEDOT:PSS hydrogels was performed using Web of Science. This search yielded 78 results. After reviewing abstracts, literature which discussed other conducting polymer formulations besides PEDOT:PSS or fabrication techniques that were not additive manufacturing were eliminated. After further reviewing, articles that reported both modulus and conductivity values of printed hydrogels were selected for presentation in Figure 3I.

### Statistical analysis

For statistical analysis, ordinary one-way ANOVA with Sıdak’s or Dunnett’s multiple comparison tests were performed with GraphPad Prism. Results are presented as means with positive and negative standard deviation. Differences were categorized as statistically significant for p < 0.05. Pearson correlation test was performed using GraphPad Prism.

## Author Contributions

S.S.O. and A.L.R. envisioned this study and wrote this manuscript. S.S.O., J.P., T.L., A.P.G., and A.L.R. were involved in data interpretation. S.S.O developed the ink, printing methods and conducted mechanical and electrical characterizations. J.P conducted XPS measurements. T.L. designed schematics and performed SEM imaging. S.S.O and A.P.G performed confocal imaging. S.S.O and R.M.A performed ink rheological characterizations. S.S.O designed and performed experiments relating to material stability in cell culture media and cell culture including quantification of seeding, viability and proliferation. B.A.S. contributed to experimental design and data interpretation for mechanical characterization. J.Y., C.O., S.M, and L.F. prepared materials and performed measurements of hydrogel dimensions including swelling over time.

## Acknowledgements

This research was supported in part by Washington University in St. Louis through the Center for Regenerative Medicine Seed Grant, the Women’s Health Technologies Collaboration Initiation, and the McDonnell Center for Cellular and Molecular Neurobiology Small Grant. The authors further acknowledge support from the National Science Foundation through FR #2319060 and the Center for Engineering MechanoBiology (CEMB), an NSF Science and Technology Center, under grant agreement CMMI #15-48571. S.S.O acknowledges support from the McDonnell International Scholars Academy. The authors acknowledge financial support from Washington University in St. Louis and the Department of Mechanical Engineering and Materials Science Shared Instrument Group for assistance from staff and use of instruments. The authors would also like to thank Dianne Duncan (Director of the Washington University in St. Louis Biology Department Imaging Facility) for technical assistance regarding imaging as well as Gregory Strout (Research Specialist) and Xue Wen Ng (Staff Scientist) at Washington University Center for Cellular Imaging for help with SEM sample preparation.

## Supplementary Information

**Supplementary Figure 1.**
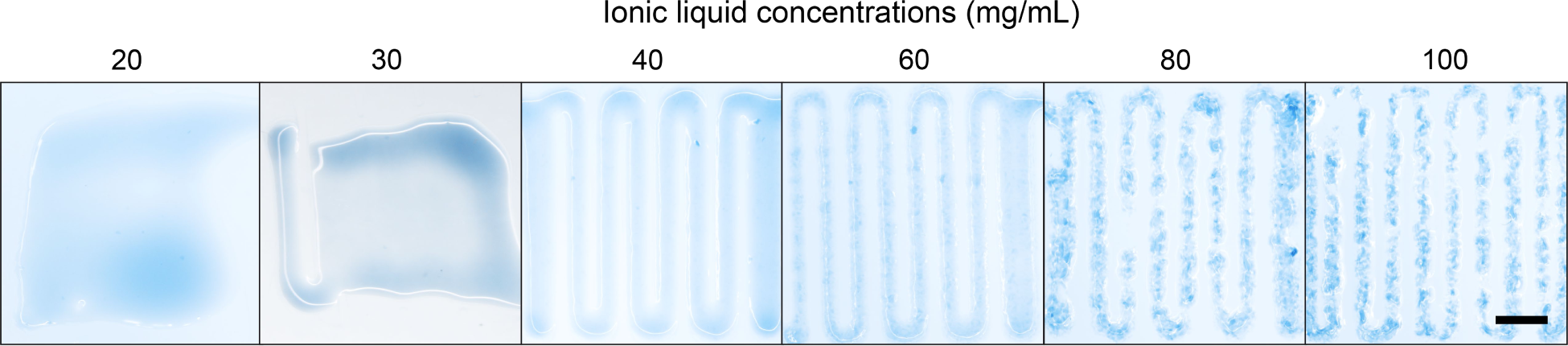
Representative images of 1 layer extrusion on glass with PEDOT:PSS-IL mixtures. Mixtures of ionic liquid (IL) concentrations of 20 and 30 mg/mL spread on substrate; 40 – 60 mg/mL resulted in continuous and defined printed lines; 80 – 100 mg/mL resulted in defined but fragmented lines. Scale bar = 2 mm.

**Supplementary Figure 2.**
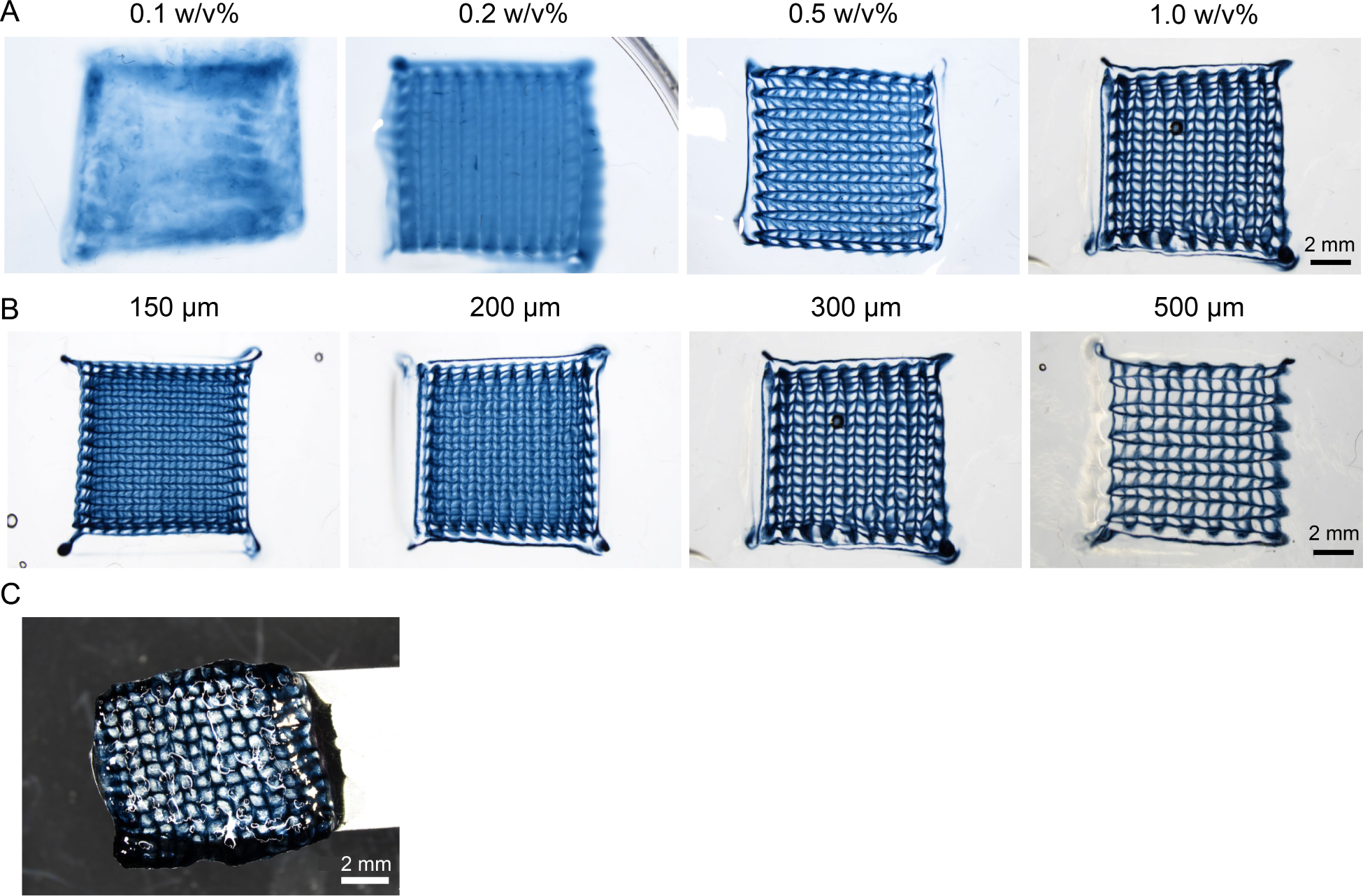
Carbopol support medium allows for printing defined structures; however, insufficient removal of support medium from pores makes it unsuitable for fabrication of PEDOT:PSS hydrogels. A) The effect of increasing concentration of Carbopol on printing. By increasing concentration of Carbopol, diffusion decreased, and more defined structures were obtained for concentrations > 0.5 w/v%. B) Printed structures with different pore sizes at 1 w/v% Carbopol concentration. At lower pore sizes (≤200 µm), although defined structures were also obtained, ink diffusion into surrounding pores was also observed. At larger pore sizes (≥ 300 µm), clearer and more distinct pores were obtained with reduced diffusion. C) Image of printed PEDOT:PSS structure after attempted washing to remove Carbopol. Carbopol was not successfully removed by washing with phosphate buffered saline (1X – 20X concentrations) as evident by the filled pores.

**Supplementary Figure 3.**
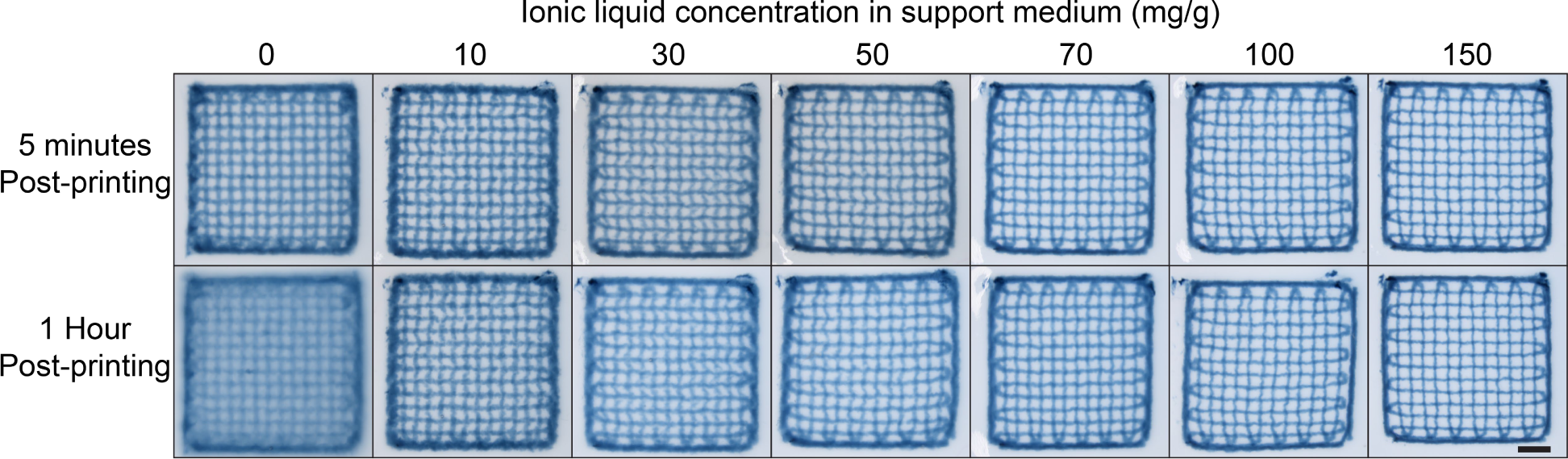
Brightfield images of PEDOT:PSS grids printed in agarose granular gel support medium with varying concentrations of ionic liquid. Scale bar = 2 mm.

**Supplementary Figure 4.**
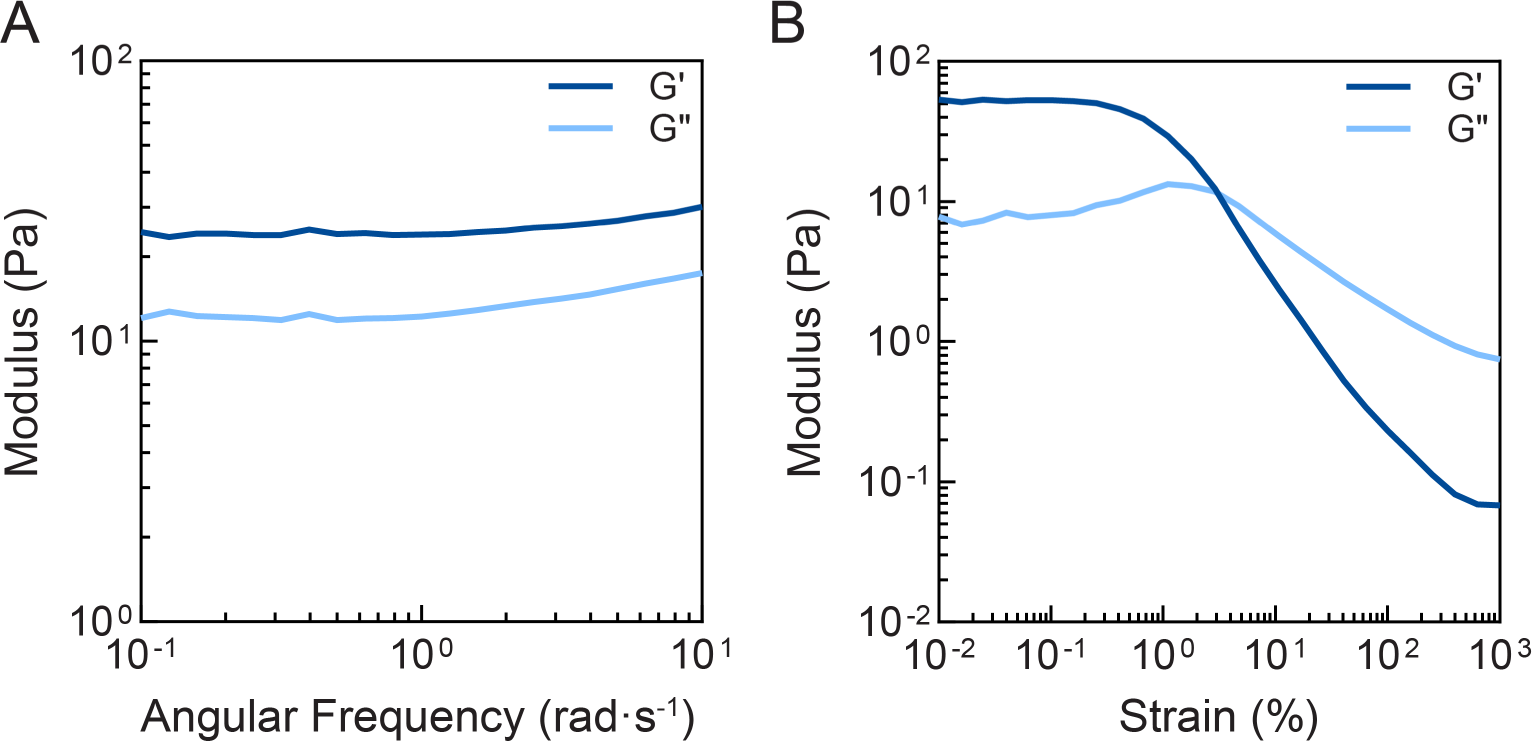
Oscillatory rheological characterization of agarose support medium without ionic liquid. (A) frequency sweep confirms expected gel viscoelasticity (G’ independent of frequency and G’>G” for all frequencies) (B) support medium yields from gel to liquid at higher strains (1 rad⋅s^-1^ angular frequency).

**Supplementary Figure 5.**
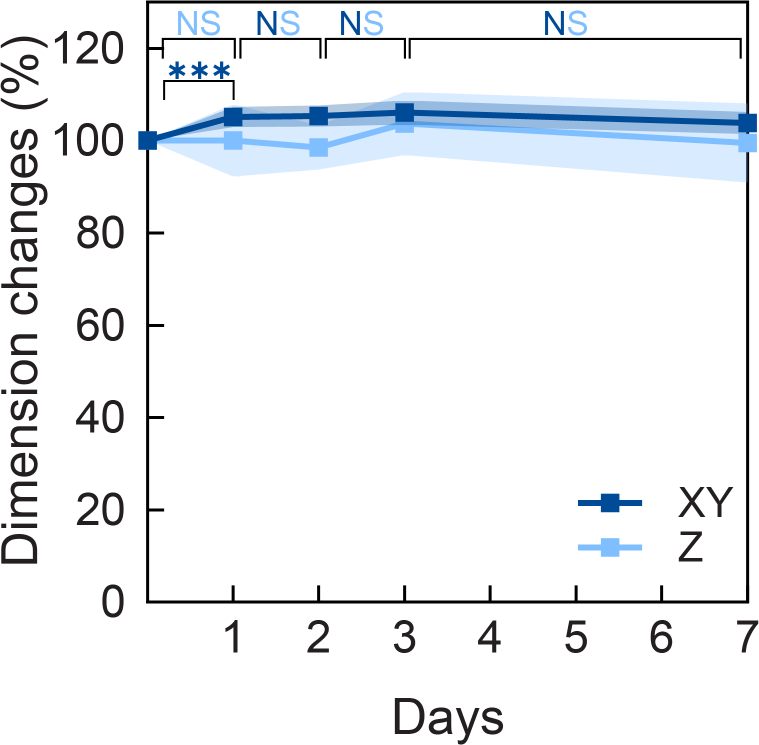
Percentage change in dimensions of PEDOT:PSS hydrogels over 7 days incubation in deionized water. Hydrogels reach equilibrium swelling within 24 hours and with no significant changes over the rest of period. Isotropic swelling was also observed (*xy*/*z* =1.05 ± 0.08). Mean and standard deviation presented, N=7. One-way analysis of variance (ANOVA) and Šidák’s multiple comparison test. ****P≤0.0001, ***P≤0.001, **P≤0.01, *P≤0.05, Non-Significant (NS) P>0.05. Alternating colors indicate that non-significance applies to both *xy* and *z* axes, while single colors are specific to the indicated axis.

**Supplementary Figure 6.**
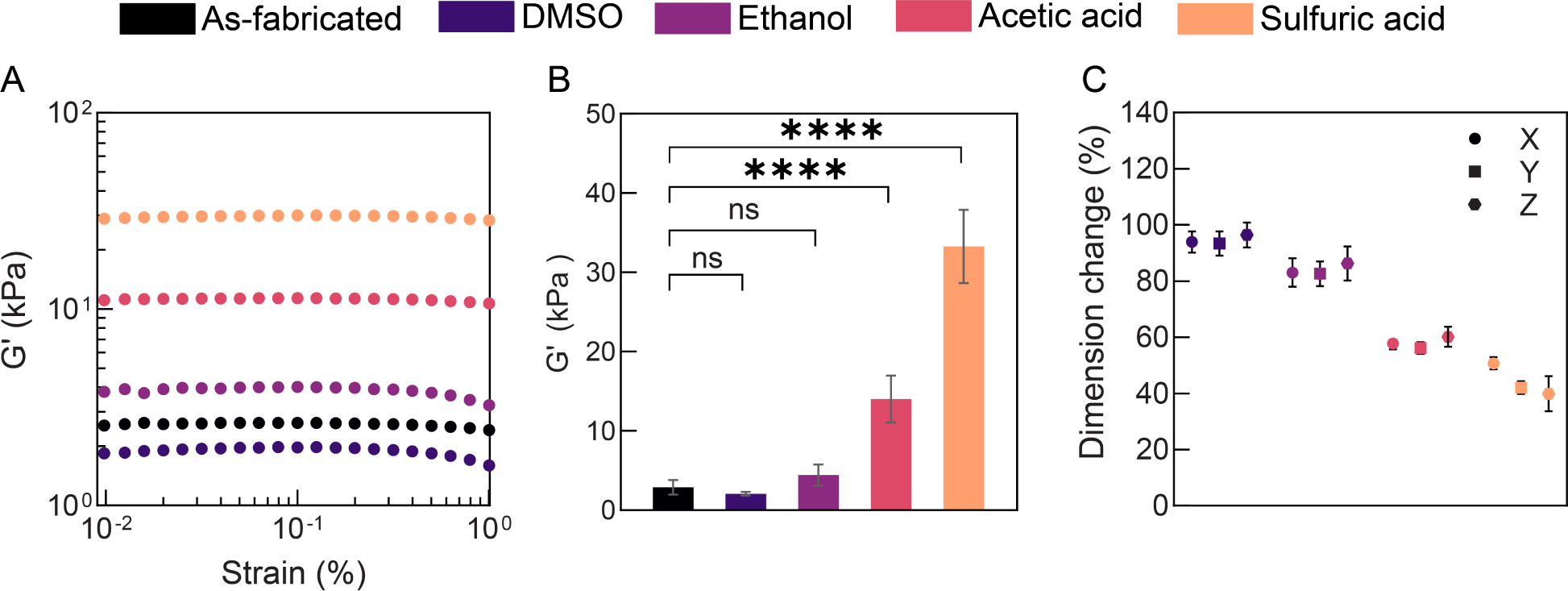
Influence of post-treatment on PEDOT:PSS hydrogel properties. A) Representative strain amplitude sweep of treated and untreated hydrogels to determine the storage modulus (10 rad⋅s^-1^, 25 °C). B) Storage modulus obtained from A. One-way analysis of variance (ANOVA) and Dunnett’s multiple comparison test performed. ****P≤0.0001, ***P≤0.001, **P≤0.01, *P≤0.05, Non-Significant (NS) P>0.05. C) Post-treatment causes a decrease in dimensions. The shrinkage occurs nearly isotopically (*xy/z* = 0.94 – 1.16). N≥4. Mean and standard deviation presented.

**Supplementary Table 1.** Table of values for material characterization of post-treated hydrogels. N≥4, mean and standard deviation presented. One-way analysis of variance (ANOVA) and Dunnett’s multiple comparison test performed. ****P≤0.0001, ***P≤0.001, **P≤0.01, *P≤0.05, Non-Significant (NS) P>0.05.

| Condition | G' (kPa) | E (kPa)<br>E= 2G'(1+ν) | Water<br>fraction (%) | σ (S/m) | PSS/PEDOT<br>ratio |
| --- | --- | --- | --- | --- | --- |
| As-fabricated | 2.90 ± 0.914 | 8.69 ± 2.74 | 99.2 ± 0.04 | 44.3 ± 14.5 | 1.74 ± 0.205 |
| DMSO | 2.07 ± 0.228<br>P = 0.9721 | 6.20 ± 0.685<br>P = 0.9721 | 99.3 ± 0.292<br>P > 0.9999 | 207 ± 10.8<br>P = 0.1432 | 1.55 ± 0.0567<br>P < 0.0001 |
| Ethanol | 4.42 ± 1.33<br>P = 0.8147 | 13.3 ± 4.00<br>P = 0.8147 | 98.7 ± 0.355<br>P = 0.9874 | 233 ± 22.6<br>P = 0.0751 | 1.57 ± 0.0801<br>P = 0.0001 |
| Acetic acid | 14.0 ± 2.96<br>P < 0.0001 | 42.1 ± 8.88<br>P < 0.0001 | 95.8 ± 1.96<br>P = 0.0850 | 946 ± 86.5<br>P < 0.0001 | 1.44 ± 0.0375<br>P < 0.0001 |
| Sulfuric acid | 33.3 ± 4.61<br>P < 0.0001 | 99.8 ± 6.92<br>P < 0.0001 | 89.7 ± 4.75<br>P < 0.0001 | 1891 ± 284<br>P < 0.0001 | 1.21 ± 0.0740<br>P < 0.0001 |

**Supplementary Figure 7.**
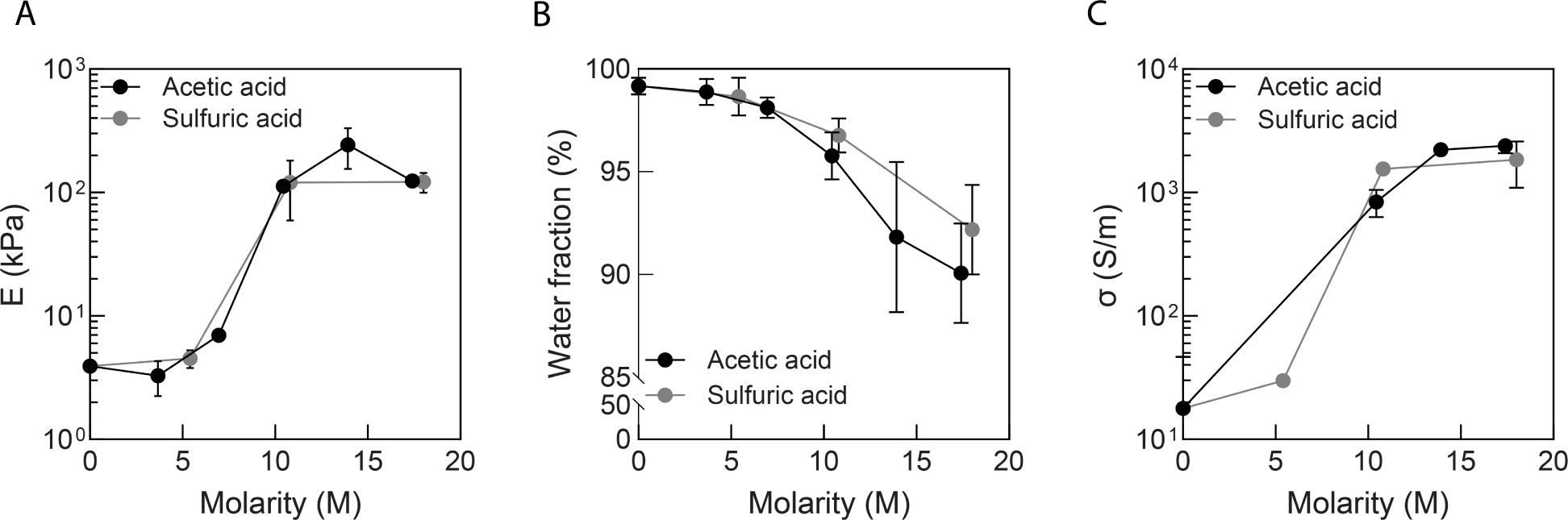
Manipulation of acid concentration changes material properties. A) elastic modulus, B) water fraction and C) conductivity. N≥2, mean and standard deviation presented. Elastic modulus obtained from compression testing of hydrogels using a TA instruments ElectroForce 3200 System equipped with a 45 N load cell. Strain rate set to 2 mm min^-1^ and samples were compressed to a 1 mm displacement. Elastic modulus was obtained from the linear region of the stress strain curve.

**Supplementary Figure 8.**
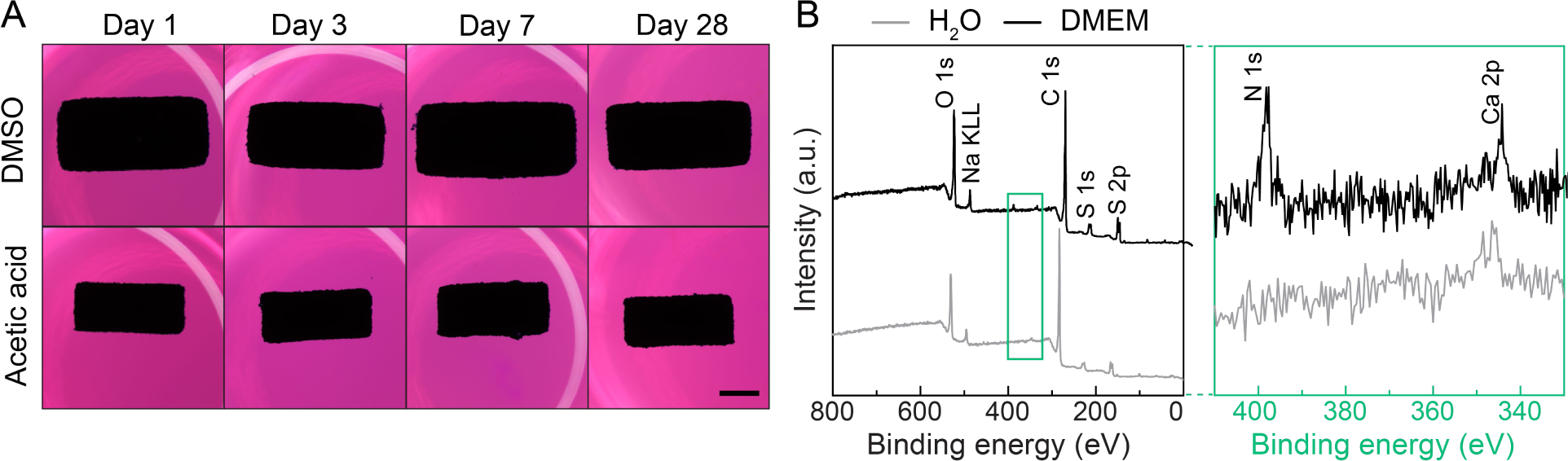
Stability analysis of PEDOT:PSS hydrogels in cell culture conditions. A). Representative images of gels post-treated with DMSO and acetic acid, scale bar = 2 mm. Hydrogels show no visible fragmentation or breakdown over 28 days for both conditions. B) DMSO treated samples before (in deionized water) and after incubating in DMEM showing presence of additional peaks. Green boxes highlight the binding energy range covering nitrogen and calcium.

**Supplementary Figure 9.**
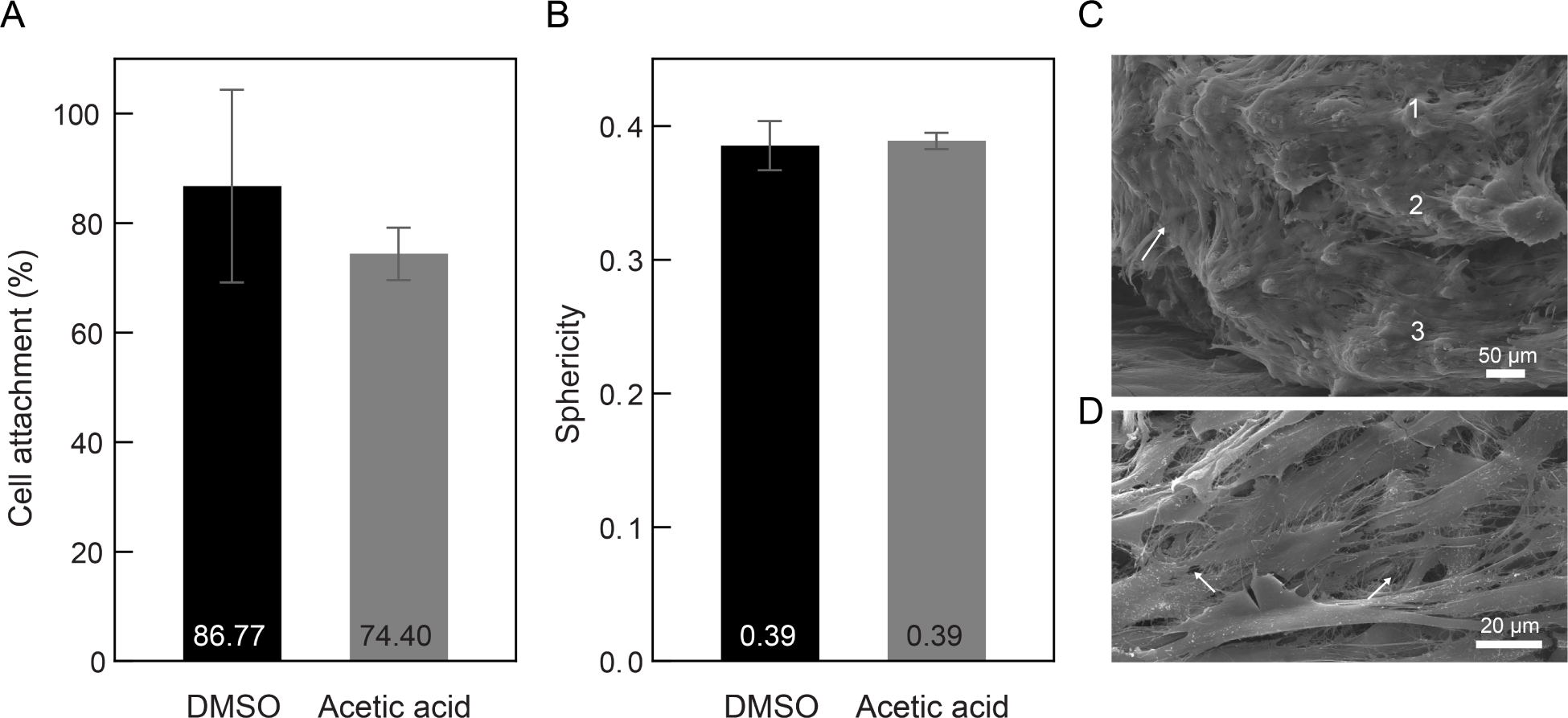
Cell-material interfacing of PEDOT:PSS hydrogel scaffolds. Quantification of (A) percent cell attachment from seeding aliquots and (B) sphericity one day after seeding. C-D) Representative SEM images of cells (C) penetrating multiple layers of the scaffold and (D) depositing ECM on acetic acid treated scaffolds after seven days. Samples for SEM were processed as described in the Methods, with the exception that EGDE crosslinker was not used. Mean and standard deviation presented. N=4.

**Supplementary Figure 10.**
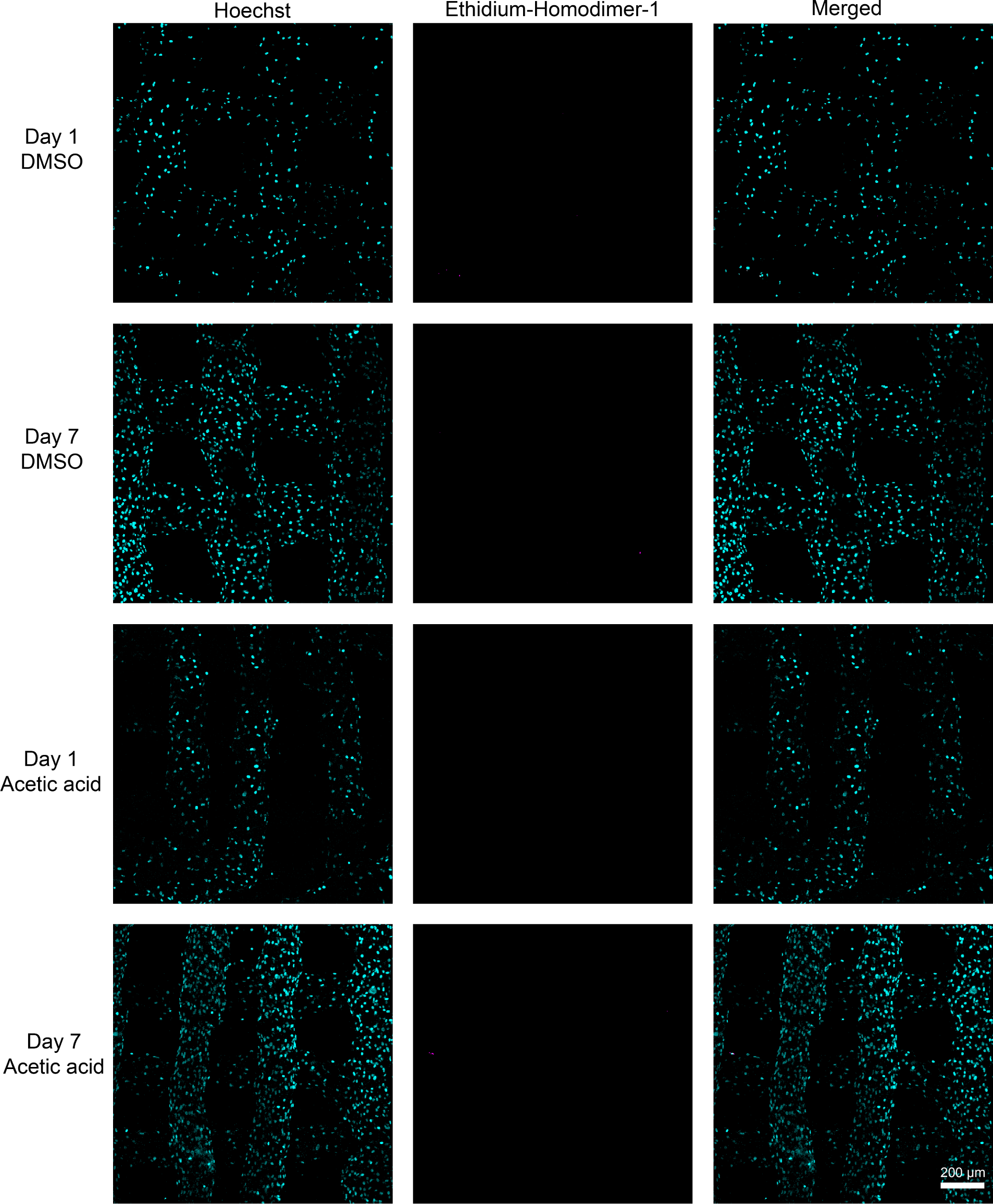
Representative fluorescence microscopy maximum intensity projections of Z stacks of human dermal fibroblasts on 3D printed PEDOT:PSS hydrogel scaffolds. Cells are stained with Hoechst and ethidium homodimer-1 for quantification of cell viability. Hoechst stains for all nuclei (cyan) and ethidium homodimer-1 stains for dead nuclei (magenta).

